# Climate change will lead to pronounced shifts in the diversity of soil microbial communities

**DOI:** 10.1101/180174

**Authors:** Joshua Ladau, Yu Shi, Xin Jing, Jin-Sheng He, Litong Chen, Xiangui Lin, Noah Fierer, Jack A Gilbert, Katherine S Pollard, Haiyan Chu

## Abstract

Soil bacteria are key to ecosystem function and maintenance of soil fertility. Leveraging associations of current geographic distributions of bacteria with historic climate, we predict that soil bacterial diversity will increase across the majority (~75%) of the Tibetan Plateau and northern North America if bacterial communities equilibrate with existing climatic conditions. This prediction is possible because the current distributions of soil bacteria have stronger correlations with climate from ~50 years ago than with current climate. This lag is likely associated with the time it takes for soil properties to adjust to changes in climate. The predicted changes are location specific and differ across bacterial taxa, including some bacteria that are predicted to have reductions in their distributions. These findings demonstrate the widespread influence that climate change will have on belowground diversity and highlight the importance of considering bacterial communities when assessing climate impacts on terrestrial ecosystems.

**IMPORTANCE:** There have been many studies highlighting how plant and animal communities lag behind climate change, causing extinction and diversity debts that will slowly be paid as communities equilibrate. By virtue of their short generation times and dispersal abilities, soil bacteria might be expected to respond to climate change quickly and to be effectively in equilibrium with current climatic conditions. We found strong evidence to the contrary in Tibet and North America. These findings could significantly improve understanding of climate impacts on soil microbial communities.

## INTRODUCTION

Climate change is disrupting almost all ecosystems on Earth, with widespread effects on plants and animals (1, 2). Continued climate shifts are predicted to exacerbate these effects. But even if climate stabilized today, disruptions to ecosystems would continue for some time. Two examples are the extinction debts of many long-lived, slowly reproducing species whose populations will dwindle in coming years due to environmental shifts that have already occurred (3, 4) and the colonization lags of species whose ranges are in the process of moving in response to climate change (5). Terrestrial bacteria play fundamental roles in the functioning of ecosystems and the maintenance of soil fertility (6, 7). However, despite the fact that soil bacterial communities and the processes they mediate are often highly sensitive to climate (8), we have limited knowledge of the effects of climate change on the regional distributions of soil bacteria (9-12).

This study investigates the spatial and temporal extent of legacy effects among soil bacteria and the consequences of equilibration of soil bacterial distributions to contemporary climate. By virtue of their short generation times and dispersal abilities, soil bacteria might be expected to respond to climate change quickly and to be effectively in equilibrium with current climatic conditions. However, legacy effects — defined here as community properties that persist after environmental change (13) — have been observed in soil microbial communities, which take up to 3 years to respond to drought and other environmental shifts (14-18). There is an indication of decadal-scale legacy effects in microbial enzyme activity as well (19). Microbial legacy effects are also known in agricultural (20) and other ecosystems (18, 19, 21). Moreover, because the distributions of soil bacteria are strongly influenced by edaphic characteristics [including soil pH and soil nutrient availability (22-24)], and because these soil properties change slowly over time, factors driving shifts in soil bacterial communities can reflect historic climate (25-29). Thus, soil bacterial communities may still be adjusting to existing climate change, and it may take years or decades for the full effects of existing climate change to become evident.

The Tibetan Plateau provides an ideal location to address these questions. It is the youngest (~2.4×10^8^ years), largest (~2.0×10^6^ km^2^), and highest (mean ~4000 m) plateau in the world. Understanding how climate change will affect Tibetan soil microbial communities is important: the plateau contains a vast soil carbon reservoir (30) that may become labile due to thawing permafrost and accelerated microbial metabolism (31, 32), and the region actively moderates climate in Asia and across the globe. Also, soil microbes on the Tibetan Plateau are exposed to particularly dry, cold conditions. Because the plateau is undergoing rapid climate change (33), many of the factors that drive the distributions of soil bacteria, particularly soil properties and plant communities, may still be equilibrating to the current climate. Thus, we anticipated that equilibration to existing climate change would induce dramatic shifts in the distributions of soil microbes in the region.

To test this hypothesis, we measured bacterial community composition in 180 non-agricultural soils from 60 locations across the plateau. We showed that the bacterial distributions in these soils were more closely associated with historic (climate from over 50 years ago) rather than contemporary climate. Using models of associations between current microbial communities and historic environmental factors, we show that diversity, community structure, and biogeographic patterns would shift substantially with equilibration to contemporary climate. To explore how generally applicable these findings are, we performed analogous analyses with 84 soil samples from across the United States and Canada. Our results suggest widespread increases in soil bacterial diversity in both regions and region-specific shifts in the distributions of individual taxa if these communities were to equilibrate to current conditions.

## RESULTS AND DISCUSSION

### Disequilibrium of bacterial communities with current climate

We profiled bacterial community structure using 16S rRNA gene amplicon sequencing from 180 soil samples across 60 locations in the Tibetan Plateau and obtained a total of 926,609 reads (median = 5,247 per sample, range = 3,016 to 9,926 per sample). These communities had 65,874 operational taxonomic units (OTUs, defined at ≥97% sequence similarity) and were dominated by nine phyla (Fig. S1, Table S1).

We obtained monthly maps of 10 climate variables across the plateau at 0.5-degree resolution from 1950 to 2012 (34). To dampen noise from short-term fluctuations, for each climate variable, we created climatologies by averaging values at our sampling locations over 10- and 20-year sliding windows. Results from the two durations were qualitatively similar, so we focus on the results from the 10-year climatologies. We refer to these climatologies by the dates that they span (e.g., 1950-1959 climatology averages climate data from 1950 to 1959). We also calculated 1-year climatologies from the year when samples were collected to account for effects of contemporary climate. We performed principal components analysis (PCA) on the climate data and found that the first three principal components (PC1, PC2, and PC3) account for 89% of variation while reducing its dimensionality, which is key for model selection in the following analyses. For most time periods, temperature is highly weighted in PC1, precipitation in PC2, and temperature range in PC3. In the following analyses, we use the projections of location-date combinations onto the first three principal components in lieu of the raw climate data, referring to these summaries of the climatologies as “climate variables”.

To assess whether the soil bacterial communities are in equilibrium with contemporary climate in the Tibetan Plateau, we built a regression model of OTU richness (number of OTUs) as a function of historical and contemporary climate variables. By performing all subsets model selection in which climate variables from different time periods compete with each other based on how well they can explain variation in OTU richness across sampling locations, we assessed the extent to which the contemporary distribution of bacterial diversity is associated with historic and contemporary climate (Methods). We also performed analogous analyses to assess correlations of contemporary and historical climate with Shannon diversity (evenness of OTUs) and with relative abundance of each bacterial family and OTU found in 40 or more soil samples. Soil bacterial distributions that are significantly correlated with the climate from several decades ago as opposed to the climate from the time of sampling would suggest that the distributions are out of equilibrium with contemporary climate. Consistent with this prediction, climate from before 1974 was important in predicting contemporary bacterial richness [Lindeman, Merenda, and Gold statistic (LMG) for 1960-1969 PC1 and 1960-1969 PC3 0.183 and 0.202, respectively; overall LMG = 0.385]. Contemporary climate variables were also important to prediction accuracy (2002-2011 PC2 and PC3 LMG = 0.415 and 0.200, respectively). Contemporary Shannon diversity is highly correlated with richness and hence is also predicted by both historic and contemporary climate. For models to predict the relative abundance of families and OTUs, the importance of climate across the decades spanning 1959 to 2012 was substantial and consistent (Fig. S2, Fig. S3). However, the frequency with which climate variables from different decades were predictive was bimodal: both historic variables from circa 1969 and contemporary variables were typically predictive of the distributions of families and OTUs, but variables from circa 1980 were rarely predictive (Fig. 1A, Fig. S4). This bimodality held quite generally across different climate variables (Fig. 1B) and also held when we analyzed climatologies for raw climate variables rather than the PCs (see below; Fig. S5). Furthermore, contemporary distributions of families and OTUs were often simultaneously associated with both historic and contemporary climate (Fig. 1C, Fig. S6). These results suggest that contemporary distributions of the diversity of soil bacteria and of individual taxa are associated with climate from close to 50 years ago and from today.

**FIG 1.**
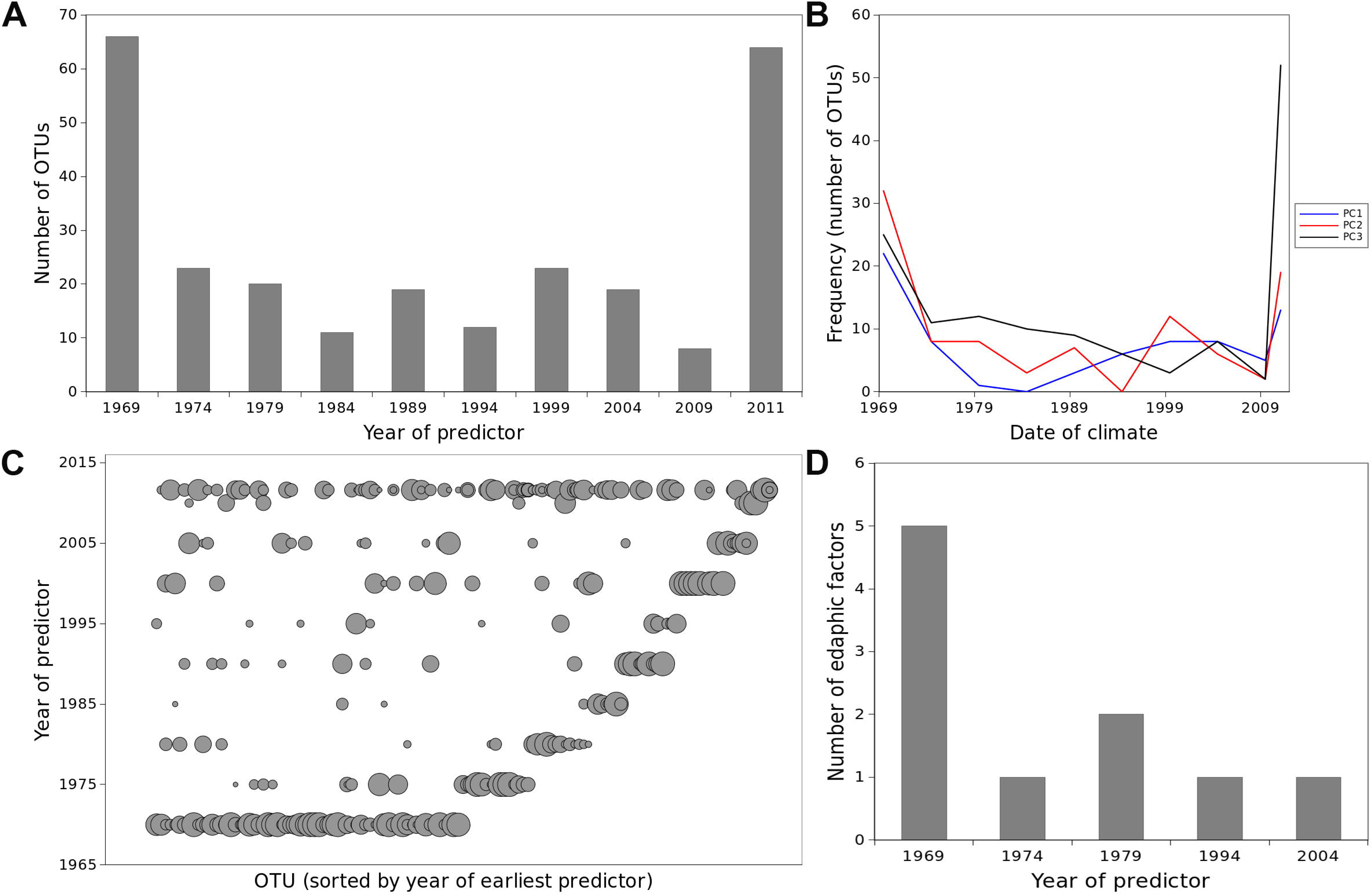
Distributions of soil bacteria in Tibet lag behind shifts in climate by up to 50 years. (A) The number of OTUs associated with climate from different years. A given OTU can be associated with climate from multiple years; the 2011 category represents climate from the year of sample collection. Lags are indicated by the association of many OTUs with climate from prior to 2011 and in many cases prior to 1980. (B) OTUs were associated with climate from both contemporary and historic values of most climate variables. (C) Most OTUs associated with historic climate were also associated with contemporary climate. Symbol size is proportional to the strength of the association, and OTUs (x-axis) are ordered by the earliest year of climate with which they were associated. (D) Soil properties were also associated with historic climate, suggesting that the lags in bacterial distributions may be mediated or associated with lags in soil properties.

Historical climate may be an important predictor of contemporary microbial distributions because soil edaphic characteristics often follow historic conditions (35). Indeed, we found that historical climate variables were more predictive than current climate of soil edaphic characteristics (Fig. 1D), and that five key soil edaphic characteristics (i.e., DON (dissolved organic nitrogen), soil organic carbon, total carbon, DOC (dissolved organic carbon) and total nitrogen) were particularly strongly correlated with climate from before 1980 (Fig. S7). Even though most soil microbes likely have short generation times, the diversity and composition of soil microbial communities appear to be strongly influenced by soil properties that change slowly over time (36). Like the microbial communities, these soil properties are out of equilibrium with contemporary climate.

These findings cannot be explained by climate cycling because most climate variables have trended consistently over this period. Moreover, the samples we analyzed are from undisturbed soils, and anthropogenic impacts other than climate change (e.g., land use change or pollution) are unlikely to have generated the disequilibria because they cannot account for the stronger correlations between current bacterial distributions and historic, rather than contemporary, climatic conditions. To evaluate the extent to which our findings might be influenced by modeling choices, we conducted two robustness analyses. First, we performed regression modeling with the original climate variables rather than projections onto principal components, using a modified model selection procedure because all subsets selection is computationally prohibitive on 120 climatologies (10 variables x 12 time periods per location; see Methods). Second, we repeated our investigation of the association between climate and bacterial distributions using gradient boosting rather than standard regression. Conclusions from both of these alternative approaches were highly concordant with our primary findings, indicating that our results are not artifacts of a particular model or way to quantify climate data.

### Widespread shifts in distributions of Tibetan soil bacteria

We next asked whether soil bacterial diversity would increase or decrease as communities equilibrated to current climatic conditions. To answer this question, we projected the models, which were fit with historical and contemporary climate data, to contemporary climate data. Specifically, by inputting 2002-2011 climatology data into the models with best performance (e.g., models utilizing 1960-1969 PC1 and PC3), we forecast how bacterial diversity and relative abundance would change if the distributions of bacteria were to equilibrate to 2002-2011 climatology. To understand the extent of these changes, we sought to answer four specific questions. (1) With equilibration, would diversity and relative abundance predominantly increase, decrease, or remain unchanged across the sampling locations? (2) How would shifts in diversity and relative abundance compare to existing spatial variation in diversity and relative abundance? (3) Would locations with currently high diversity or relative abundance experience different changes compared to the locations with low diversity and low relative abundance (i.e., “rich get richer” versus homogenization)? (4) Would intersample variability increase or decrease in the future?

We forecast that (1) richness and Shannon diversity would increase across 75% and 72.9% of the sampling locations, respectively, with an average magnitude of +7.5% (standard error 1.5%) for richness and +2.1% (standard error 0.4%) for Shannon diversity (Fig. 2A). We further forecast (2) that shifts in diversity within samples would be of similar magnitude to existing inter-sample differences in diversity, suggesting major, although not unprecedented, shifts in diversity (Fig. S8, Fig. S9). We forecast that (3) locations with low diversity would experience the largest increases in diversity, and locations with high diversity would experience little or no increases in diversity (Fig. 2B, Fig. S10). This latter trend might suggest that intersample variation in diversity levels would decrease in the future, but our forecasts indicated (4) intersample variability would remain relatively constant (Fig. S8, Fig. S9). These forecasts suggest major changes in the spatial distribution of diversity in soil microbial communities across the Tibetan Plateau with equilibration to existing climate change.

**FIG 2.**
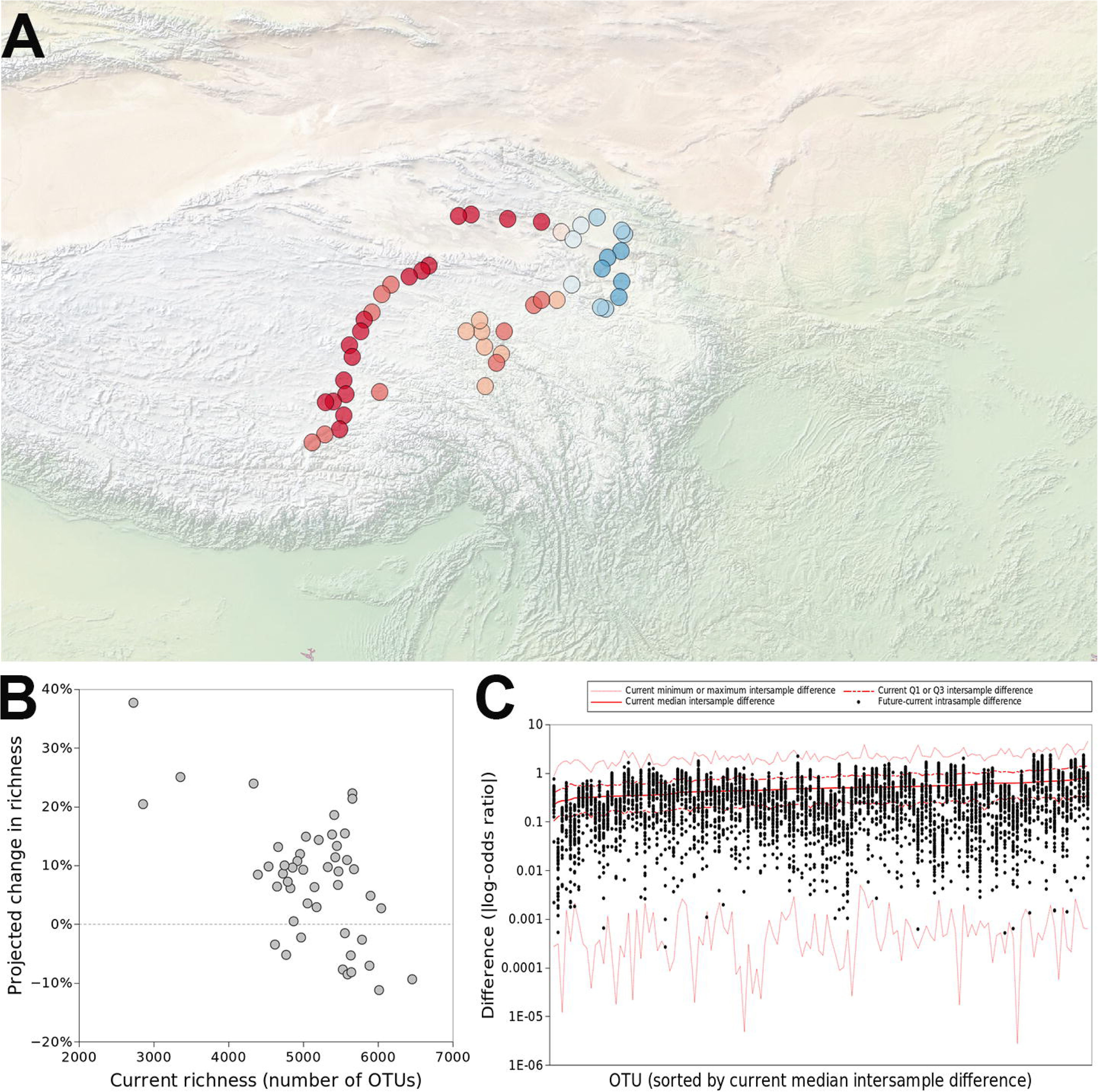
With equilibration to contemporary climate, the distributions of soil bacteria in Tibet would shift substantially. (A) Across most of the locations sampled, richness would increase, although in some locations it would decrease. Red and blue indicate increases and decreases in richness, respectively. (B) Increases in richness would be greatest in locations that have relatively low richness; locations with higher contemporary richness would see little change, or even decreases in richness. (C) The magnitude of shifts in relative abundance of OTUs with equilibration would be comparable to contemporary intersample variability in their relative abundance. Red lines indicate current intrasample differences in relative abundance; black dots represent the projected shifts in relative abundance with equilibration.

Further supporting this contention, we used the regression models developed above to forecast how bacterial richness would shift if bacterial distributions were to equilibrate across the Tibetan Plateau. Specifically, we projected maps of current richness using the historic and contemporary climate conditions that were selected for the models. Using just contemporary climate conditions, we then projected richness maps if bacterial distributions were to equilibrate to contemporary climate. These projections (Fig. 3) were consistent with foregoing results, suggesting widespread increases in richness across Tibetan Plateau.

**FIG 3.**
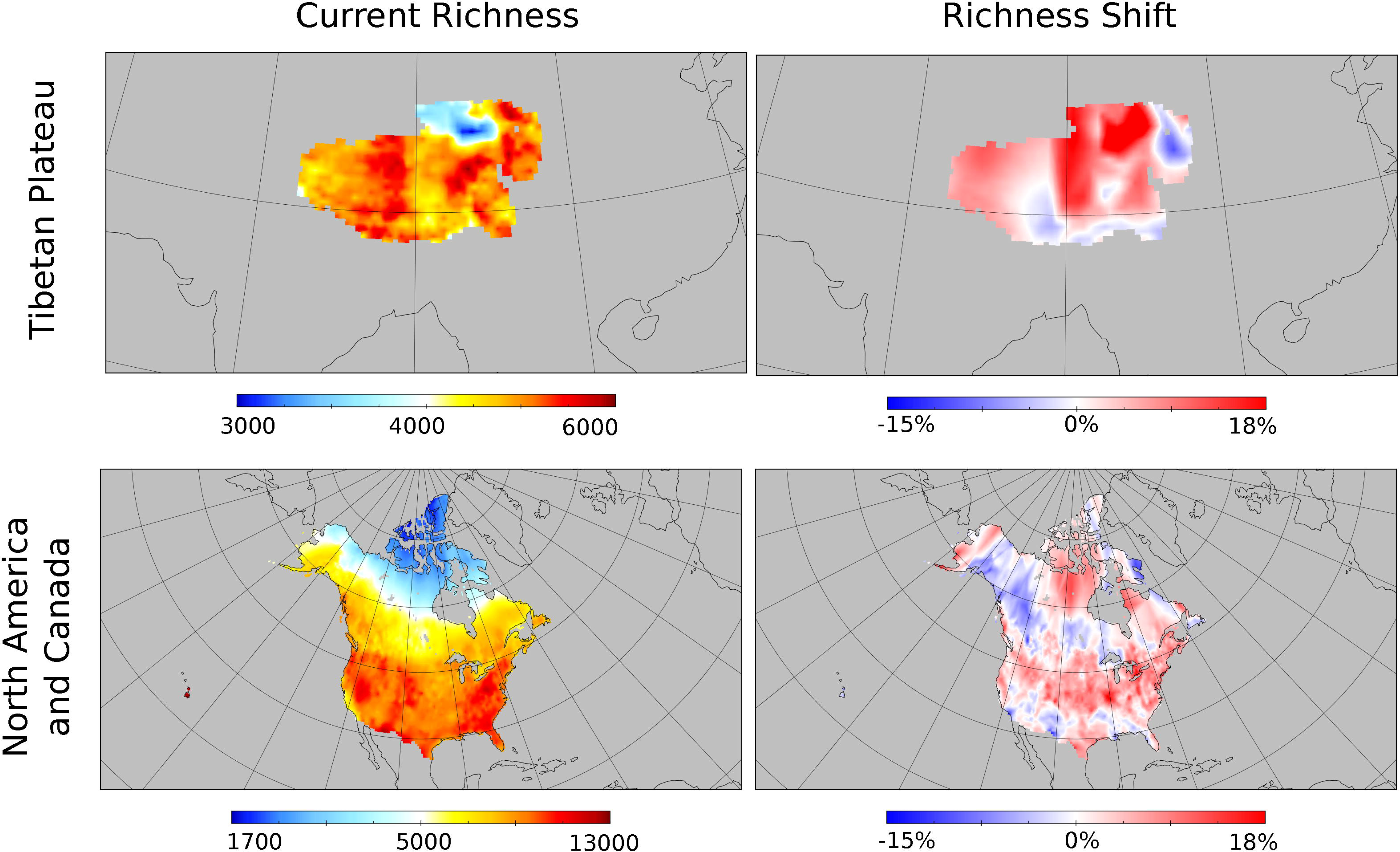
As in the Tibetan Plateau, in North America the distributions of soil bacteria lag behind climate change. If they were to equilibrate to contemporary climate, then substantial shifts in their distributions would result. (A) Climate variables from prior to 1980 are predictive of the distributions of many OTUs; the 2005 category represents the climate from the time when samples were collected. (B) If the distribution of an OTU is associated with historic climate variable, then it is also likely to be associated with contemporary climate. Symbol size is proportional to predictive power. (C) With equilibration to contemporary climate, the shifts in relative abundance would be of similar magnitude to existing differences in relative abundance between microbial communities across northern North America. Red delineates existing intersample variability in relative abundance; black dots indicate projected intrasample shifts.

Turning to how the relative abundance of individual taxa would respond to equilibration to current climate, we found (1) that different taxa would respond hetergeneously: some would increase across all sampling locations, but others would decrease across all sampling locations; the magnitude of changes ranged from non-significant to over 100% for different taxa (Table S2). For example, Mycobacteriaceae and Rubrobacteraceae (families of Actinobacteria), Bacillaceae (a family of Firmicutes), Rhodobiaceae and Rhizobiaceae (families of Alphaproteobacteria) are predicted to increase consistently in most locations of the plateau, while Flavobacteriaceae (a family of Bacteroidetes), Nakamurellaceae, and Nocardiaceae (families of Actinobacteria) are predicted to decrease in the majority of locations. Overall, there was no consistent trend across all families or OTUs (e.g., most families increasing). However, (2) shifts in relative abundance across taxa would consistently be of similar magnitude to existing intersample differences in relative abundance (Fig. 2C, Figs. S11-S13). That is, the projected changes in community composition are on par with existing inter-site variation in communities across the sampled locations. Furthermore, (3) locations with low relative abundance would experience larger changes than locations with high relative abundance (Fig. S14, Fig. S15). Finally, although the latter shifts could act to even out the spatial distribution of relative abundance, this does not appear to be the case: (4) with equilibration, intersample differences in relative abundance would be similar to contemporary intersample differences (Fig. S16, Fig. S17). Thus, across the Tibetan Plateau, different taxa would undergo varying shifts in relative abundance with equilibration, with overall widespread, substantial impacts to communities.

Our forecasts of shifts with equilibration to existing climate assume temporal niche conservatism, which means that bacteria are associated similarly with environmental conditions over time, as they equilibrate to changes. Bacteria could shift to occupy different niches in the time that it would take their distributions to equilibrate to contemporary climate. However, incorporating such shifts into our models would introduce substantial complexity and numerous assumptions. Thus, our forecasts can be taken as baseline estimates: future analyses that incorporate additional complexity may add to these results.

### Increases in diversity of northern North American bacteria

To assess whether the predicted responses of Tibetan Plateau soils to climate change are similar to those for other regions of Earth and at larger spatial scales, we performed similar analyses using historic climate data and published soil microbial community data from 84 locations across northern North America (Fig. S18) (24, 37). Strikingly, several of the major trends from the Tibetan Plateau were also observed in northern North America. First, historical climate variables were strong predictors of taxon abundance and diversity metrics. For instance, soil bacterial richness was predicted by 1960-1969 climatologies in both regions (in northern North America, LMG for 1960-1969 PC2 and PC3 0.057 and 0.227, respectively, with LMG for PC1 in 1975-1984 = 0.716). Second, across families and genera, relative abundances were commonly predicted by both historical and contemporary climate in northern North America (Fig. 4A and 4B, Fig. S19, Fig. S20). When they were included via model selection, historical climate variables were important predictors (Fig. S21, Fig. S22). Furthermore, historical values of most climate variables were predictive (Fig. S23, Fig. S24). Finally, when these models were projected to contemporary climate, the forecast outcomes of equilibration were similar to those in the Tibetan Plateau: richness and Shannon diversity would increase across 76.0% and 73.0% of samples, respectively (Fig. S25). Projecting maps of the increases in richness, these increases in richness would be geographically widespread (Fig. 3), and shifts in diversity and relative abundance would be of similar magnitude to existing spatial variation in these quantities (Fig. 4C, Figs. S26-30).

**FIG 4.**
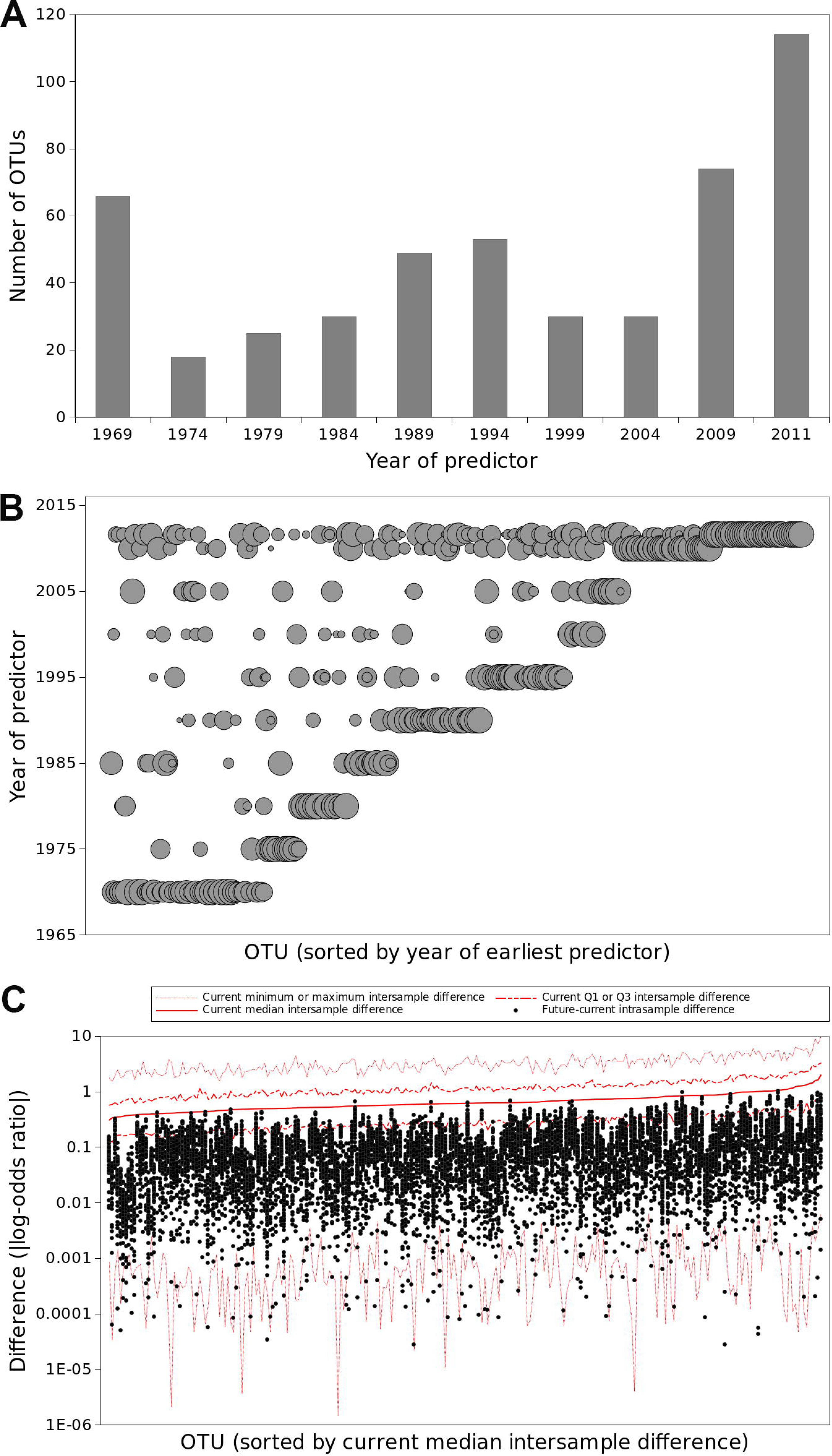
Associations between taxa and climate over time. (A) The number of OTUs associated with climate from different years in North America. (B) Most OTUs associated with historic climate were also associated with contemporary climate. Symbol size is proportional to the strength of the association, and OTUs (x-axis) are ordered by the earliest year of climate with which they were associated. (C) The magnitude of shifts in relative abundance of OTUs with equilibration would be comparable to contemporary intersample variability in their relative abundance. Red lines indicate current intrasample differences in relative abundance; black dots represent the projected shifts in relative abundance with equilibration.

Despite these similarities, we observed several important differences between our models for northern North America and the Tibetan Plateau. In northern North America, changes in richness after equilibration to current climate would be uncorrelated with current richness (Fig. S31), although changes in Shannon diversity would be negatively associated with current Shannon diversity (Fig. S32). Furthermore, families and OTUs would generally have the greatest changes in relative abundance in locations where they are currently rare (Fig. S33, Fig. S34), although the distribution of future intersample differences in relative abundance is similar to current intersample differences (Fig. S35, Fig. S36). The most striking difference between the regions is that individual taxa have very different forecast changes in their distributions (R^2^=0.053 for correlation, between regions, of fraction of locations where families would increase). For example, Beijerinckiaceae (a family of Alphaproteobacteria) and Acidobacteriaceae (a family of Acidobacteria) are predicted to increase in the relative abundance in most locations of northern North America (Table S3), while Methylobacteriaceae (a family of Alphaproteobacteria) and Cellulomonadaceae (a family of Actinobacteria) are predicted to decrease. Thus, our results demonstrate different responses among specific bacterial taxa and between Tibetan Plateau and North America despite parallel trends towards higher diversity.

### Proximate causes of the disequilibrium

The disequilibrium between bacterial distributions and contemporary climate is likely due to the soil properties being out of equilibrium with contemporary climate. To explore this hypothesis, we analyzed correlations of current soil properties with patterns of bacterial diversity at the locations that we sampled in Tibetan Plateau. Soil factors, including DON and DTN, were significantly correlated with bacterial OTU richness, and DON, SOC, TC and DTN were significantly correlated with bacterial community structure (Tables S4 and S5). In some other studies at this scale, pH had a strong association with soil bacterial richness (38), particularly in acidic soils (39) and when a wide range of pH values is observed. However, at sampling locations in this study, the soil carbon-to-nitrogen ratio (C:N) was more important, being negatively correlated with richness (r^2^ = 0. 26, P < 0.001; Fig. 5, and Table S4) and community structure (Bray-Curtis dissimilarity; r =0.44, P=0.001; Fig. 6 and Table S5). C:N is also the best predictor of the relative abundance of some, but not all, individual taxa. The high altitude and low temperatures on the plateau reduce C degradation rates and lead to N limitation (30), resulting in elevated C:N ratios and high inorganic C in dry areas. Soil moisture, which correlated with C:N, showed similar associations with richness and community composition (Fig. S37, Table S4). The relative abundance of specific taxa have both negative (e.g., *Alphaproteobacteria*) and positive (e.g., *Bacteroidetes*) correlations with C:N ratio and soil moisture (Fig. S38, Fig. S39). We found that C:N ratios are more closely associated with historical rather than contemporary climate (climatology of strongest association: 1960-1969), suggesting a mechanism through which bacterial distributions are out of equilibrium with contemporary climate: distributions of soil properties lag behind shifts in climate, which in turn cause the distributions of bacteria to lag. However, we cannot rule out the possibility that other factors, such as lagging distributions of vegetation, could be a driver of the lagged relationship of bacteria to climate, or that bacteria are inherently slow to respond to climate change, irrespective of changes to soil and plants.

**FIG 5.**
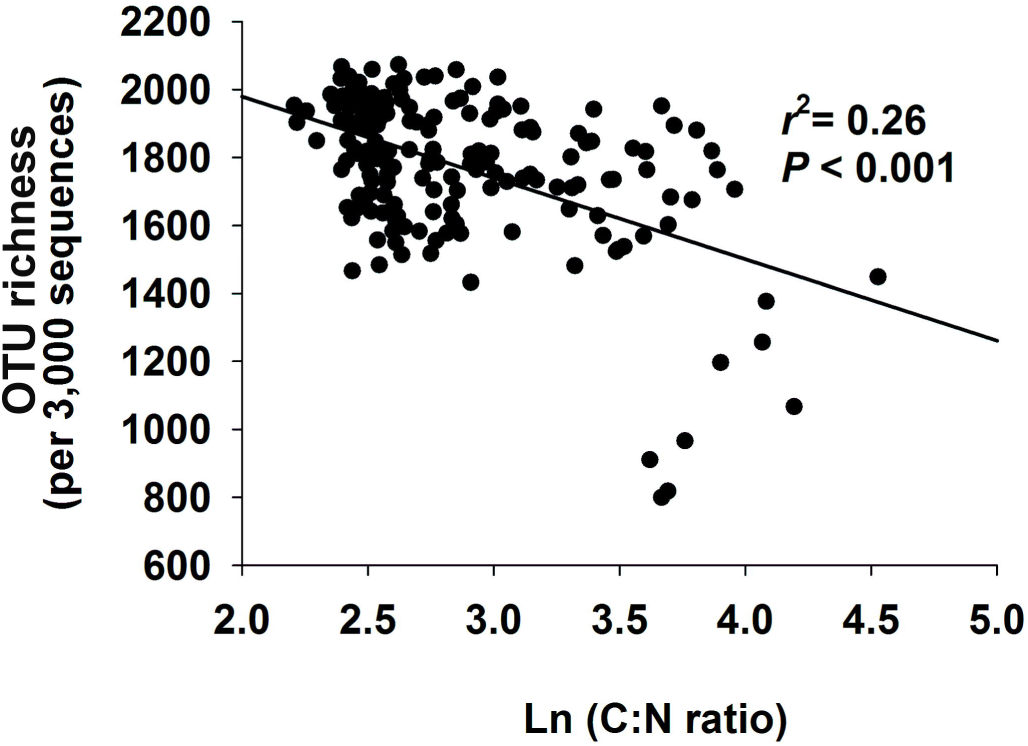
Relationship between bacterial OTU richness and soil C:N ratios.

**FIG 6.**
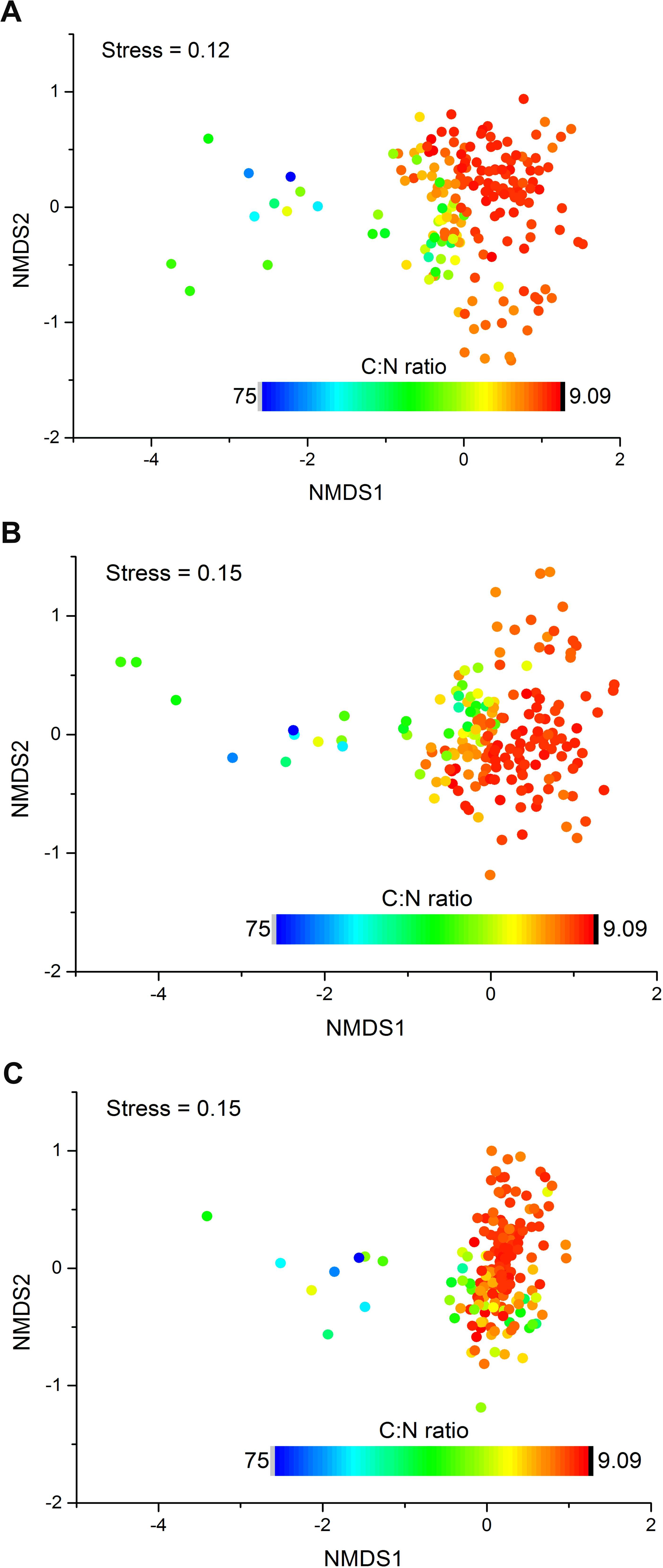
Bacterial communities in Tibetan Plateau soils are associated strongly with soil C:N ratios. Bacterial community compositional structure in the Tibetan Plateau soils as indicated by non-metric multidimensional scaling plots. Sites are color coded according to soil C:N ratios. A: Based on Bray-Curtis distance. B: Based on Unweighted Unifrac distance. C: Based on Weighted Unifrac distance.

### Conclusions

Soil bacteria appear to follow soil characteristics in showing a significant lagged response to a changing climate across many decades, a pattern evident across both the Tibetan Plateau and northern North American. If bacteria could equilibrate to existing climate change, widespread increases in bacterial diversity and shifts in community composition would occur. Similar to the diversity debts and colonization lags of macroorganisms (3, 5), further climate change is likely to exacerbate these changes in soil communities.

## MATERIALS AND METHODS

### Sample collection (Tibetan Plateau samples)

To survey current bacterial distributions across Tibetan Plateau, we collected 180 soil samples from 60 sites throughout Tibetan Plateau during the growing season (July to September) of 2011. At each site, we sampled three plots 40 meters apart, and collected 5–7 cores per plot at a depth of 0–5 cm, which were subsequently combined. Our sampling locations covered more than 1,000,000 km^2^ (Fig. S40, Table S6) and all of the major climate zones and grassland types across the Tibetan Plateau (Table S6). All soil samples were delivered by cooler equipped with ice packs (4°C) to the laboratory as quickly as possible, where they were stored at −20°C until processing. In addition, all vegetation in three plots (1×1m^2^ or 0.5 × 0.5 m^2^) 10 meters away from the soil sampling-plot were censored and harvested to measure aboveground biomass. At each site, one soil pit was excavated to collect samples for analyses of bulk density. From this pit, three replicate soil samples were collected at depth of 0–5 cm. Bulk density was obtained using a standard container with 100 cm^3^ (50.46 mm in diameter and 50 mm in height) and measured to the nearest 0.1 g.

### Soil characteristics (Tibetan Plateau samples)

Soil samples for C and N analyses were air-dried, sieved (2-mm mesh), handpicked to remove fine roots, and ground. Total soil C and N contents for each plot were determined by combustion (2400 II CHNS/0 Elemental Analyzer, Perkin-Elmer, Boston, MA, USA). Soil moisture was measured gravimetrically after a 10-h desiccation at 105 °C. Soil pH was determined separately on each plot at each site with a fresh soil to water ratio of 1:5 by pH monitor (Thermo Orion-868). Bulk density was calculated as the ratio of the oven-dry soil mass to the container volume. Dissolved organic carbon, dissolved total nitrogen (DTN), ammonium nitrogen (NH_4_^+^-N), nitrate nitrogen (NO_3_^-^-N) were determined as described (40).

### Molecular analyses (Tibetan Plateau samples)

Total nucleic acids from each plot were extracted from 0.5 g of soil using a FastDNA^®^ Spin kit (Bio 101, Carlsbad, CA, USA), according to the manufacturer’s instructions, and stored at −40°C. Extracted DNA was diluted to approximately 25 ng/μl with distilled water and stored at −20°C until use. A 2-μl diluted DNA sample of each plot was used as template for amplification. The V4–V5 hyper-variable regions of bacterial 16S rRNAs (*Escherichia coli* positions 515–907) were amplified using the primer set: F515: GTGCCAGCMGCCGCGG with the Roche 454 ‘A’ pyrosequencing adapter and a unique 7-bp barcode sequence, and primer R907: CCGTCAATTCMTTTRAGTTT with the Roche 454 ‘B’ sequencing adapter at the 5′-end of each primer, respectively. Each sample was amplified in triplicate with a 50 μl reaction under the following conditions: 30 cycles of denaturation at 94°C for 30 s, annealing at 55°C for 30 s, and extension at 72°C for 30 s; with a final extension at 72°C for 10 min. PCR products from each sample were pooled together and purified with an agarose gel DNA purification kit (TaKaRa) and combined in equimolar ratios in a single tube and run on a Roche FLX454 pyrosequencing machine (Roche Diagnostics Corporation, Branford, CT), producing reads from the forward direction F515.

### Bioinformatics (Tibetan Plateau samples)

Only sequences >200-bp long with an average quality score >25 and no ambiguous characters were included in the analyses (41). Filtering of the sequences to remove erroneous operational taxonomic units (OTUs) due to sequence errors and chimeras was conducted using the USEARCH tool in QIIME (42), version 1.9.0. Bacterial phylotypes were identified using uclust (43) and assigned to operational taxonomic units (OTUs) based on 97% similarity. A representative sequence was chosen from each phylotype by selecting the most highly connected sequence (44). All representative sequences were aligned by PyNAST (45). Taxonomic identity of each phylotype was determined using the Greengenes database (http://greengenes.lbl.gov). To correct for survey effort, we used a randomly selected subset of 3000 sequences per sample. In addition, details of sample collection and bioinformatics for northern North American bacteria are given in references (24, 37).

### Statistical analyses (Tibetan Plateau samples)

Correlations between diversity estimates and soil characteristics were conducted by SPSS 20.0 for windows. Non-metric multidimensional scaling analyses were performed using vegan of R 2.3.0 (44), based on dissimilarity calculated using the Bray-Curtis index, and environmental factors were fitted using the envfit and vif of vegan package. Mantel test were carried out on Bray-Curtis distances for the community (rarefaction depth 3000 sequences); Euclidean distances for the geographical coordinates scaled environmental variables within vegan.

### Historical climate data

For assessing whether historical or current climate is more predictive of current bacterial distributions, we utilized global maps of monthly historical climate records from the 0.5 degree gridded CRU TS3.21 dataset (34). The CRU TS3.21 dataset spans 1901 to 2014, but we used only records postdating 1950, because in Tibetan Plateau and North America, records prior to then are based on substantially more interpolation (46). We considered the following climate variables: frost day frequency, potential evapo-transpiration, daily mean temperature, monthly average daily minimum temperature, monthly average daily maximum temperature, vapor pressure, wet day frequency, cloud cover, diurnal temperature range, and precipitation (Table S7). We considered 10- and 20-year climatologies (i.e., summaries over one or two decades) for each of these variables as predictors, but use of both climatologies yielded qualitatively similar results so we focus on the 10-year climatology results. To test associations with contemporary climate, we used the average conditions from the year when samples were collected (2011 in Tibet and 2005 in northern North America). Inclusion of even more time specific climate data (e.g., from the month of sample collection) did not improve model performance. We performed principal components analysis (PCA) on these variables across all climatologies and locations separately in Tibet and North America. In this PCA, each location-time period combination is an observation and each of the 10 climate measurements is a variable. We performed subsequent analyses using the projections of the location-time period observations onto the first three principal component axes, which accounted for >85% of the variation.

### Modeling

To assess associations with contemporary and historic climate, we obtained from the aforementioned maps climate records for each sampling location (Supplementary Tables 8 and 9). We obtained these records for each climate variable for each 10-year climatology ending on December 31 of 2004, 1999, 1994, 1989, 1984, 1979, 1974, 1969, 1964, and 1959. For Tibet, we also obtained records for the climatology ending on December 31 of 2009 (Table S8). In addition, for both Tibet and northern North America, we obtained climate data—averaged over January 1 to December 31—for the year of sampling.

To assess associations between contemporary bacterial distributions, and contemporary and historic climate, we constructed regression models. We constructed separate models for the distributions of bacterial OTU richness) and Shannon Diversity, and the relative abundance of all families and OTUs occurring in 40 or more samples. We used leave-one-out cross validation to assess model performance and perform model selection. We performed all subsets models selection with all of the climatology dates. In the absence of any clear non-linearity, we employed linear models to further minimize the risk of over-fitting. Diversity response variables (richness and Shannon Diversity) were log-transformed prior to modeling, and relative abundance response variables were logit-transformed. To assess robustness of our findings to modeling choices, we (i) repeated regression modeling with the original climatologies rather than PCs using a two-step variable selection procedure in which the top ~5 variables were chosen using only 1960-1969 and contemporary climatologies and then all subsets model selection was performed over all time periods for these top variables (all subsets is computationally impractical with 120 variables) and (ii) fit gradient boosted regression models with all PCs rather than performing all subsets model selection.

To predict how microbial communities would shift if they were to equilibrate to contemporary climate, we substituted contemporary climate data into the models selected above, many of which used climate data from prior to 1980. We used 10-year climatologies for these substitutions. To estimate shifts in diversity and relative abundance, we took the difference between future predictions and contemporary predictions (as opposed to the difference between future predictions and contemporary observations); this procedure avoided spurious correlations that can arise from the non-zero covariance that always exists between residuals and observed values. We used Multivariate Environmental Similarity Surface [MESS (47)] to ensure that the maps of diversity that we projected did not require excessive extrapolation (Fig. S41). Our code is available at: https://github.com/jladau/SpeciesDistributionModeling.

## Data availability

The 454 pyrosequencing dataset of Tibetan soil bacteria are deposited in the DDBJ Sequence Read Archive (http://trace.ddbj.nig.ac.jp/DRASearch) with accession number: DRA001226.

## Acknowledgements

We thank Ke Zhao, Xiaoxia Yang, Congcong Shen, Huaibo Sun and Xingjia Xiang for assistance in soil sampling and lab analyses. We also thank Huayong Zhang and Jun Zeng for assistance in data analysis. This work was supported by the Strategic Priority Research Program (XDB15010101, XDA05050404) of the Chinese Academy of Sciences, the National Program on Key Basic Research Project (2014CB954002, 2014CB954004), and the National Natural Science Foundation of China (41701298, 41371254), the “135” Plan and Frontiers Projects of Institute of Soil Science (ISSASIP1641) and the National Science and technology foundation project (2015FY110100). JAG was supported by the U.S. Dept. of Energy under Contract DE-AC02-06CH11357. NF was supported by a grant from the National Science Foundation (DEB-0953331). KSP and JL were supported by the National Science Foundation (DMS-1069303), the Gordon and Betty Moore Foundation (grant #3300), the Gladstone Institutes, and a gift from the San Simeon Fund.

